# Bending stiffness of *Candida albicans* hyphae reflects adaptive behavior of the fungal cell wall

**DOI:** 10.1101/2022.03.22.485357

**Authors:** Elodie Couttenier, Sophie Bachellier-Bassi, Christophe d’Enfert, Catherine Villard

## Abstract

The cell wall is a key component of fungi. It constitutes a stiff shell which counteracts internal cell turgor pressure. Its mechanical properties thus contribute to define cell morphology. Measurements of the elastic moduli of the fungal cell wall have been carried out in many species including *Candida albicans*, a major human opportunistic pathogen. They mainly relied on atomic force microscopy, and mostly considered the yeast form. We developed a parallelized pressure-actuated microfluidic device to measure the bending stiffness of hyphae. We found that the cell wall stiffness lies in the MPa range. We the used three different ways to disrupt cell wall physiology: inhibition of beta-glucan synthesis, a key component of the inner cell wall; application of an hyperosmotic shock triggering a sudden decrease of the hyphal diameter; deletion of two genes encoding GPI-modified cell wall proteins resulting in reduced cell wall thickness. The bending stiffness values were affected to different extents by these environmental stresses or genetic modifications. Overall, our results support the elastic nature of the cell wall and its ability to remodel at the scale of the entire hypha over minutes.

## Introduction

*Candida albicans* is one of the most important fungi from a clinical standpoint. It has been estimated that 70% of healthy individuals carry *C. albicans* at any given time during their lifetime[1]. *C. albicans* is also one of the most important human opportunistic fungal pathogens. It is responsible for mucosal diseases in otherwise healthy individuals (following for example an antibiotic treatment), as well as deep-seated infections in immunocompromised patients. Disseminated candidiasis, i.e. fungal invasion of internal organs through bloodstream infections is associated with a high mortality rate (∼50%) even with current antifungal treatments[2].

The pathogenicity of *C. albicans* relies largely on its ability to grow according to distinct morphological forms ranging from unicellular budding yeasts to elongated invasive filaments called hyphae [3, 4]. A hypha consists of mononucleated compartments separated by walls (septa) which do not alter the regular tubular diameter of the overall filament (contrarily to pseudo-hyphae[5]). The filamentous forms achieve tissue penetration and invasion, whereas the yeast form promotes dissemination, acting as seeds for future hyphal colonization in local or distant organs[6]. Filamentation can be induced *in vitro* by tuning the physicochemical growth parameters such as pH or temperature, or by adding e.g. serum or N-acetylglucosamine to the culture medium [3, 7].

As pollen tubes or root hairs, the growth of hyphae is driven by an internal turgor pressure [8] combined to a directional vesicular trafficking supplying proteins and lipids to the hyphal tip. The turgor pressure is counteracted by the existence of a bilayered cell wall, composed of an outer layer of mannan and glycoproteins, and of a denser inner layer containing polysaccharides, mainly *β*-glucans and chitin [9, 10]. Any disturbance of cell wall composition and structure may impair cell physiology, and eventually cell integrity. The cell wall can be modeled as an elastic shell of decreased elasticity away from the tip[11, 12]. Refinements of such models may include viscous aspects [13]. The combination of a stiff cell wall in subapical regions and of a softer tip leads to the lengthwise growth of hyphae.

The protrusive forces developed by the forward movement of the hyphal tip reach a few *μ*N [14], and allow penetration of viscoelastic environments of rigidities up to 200kPa[15], which is also the upper limit of the range of soft tissues rigidities [16]. To achieve penetration, hyphal adhesion is instrumental[17]. In situations where the resistive forces experienced by the hyphal tip exceeds the adhesive forces, hyphae must also resist buckling[15, 18]. The beam theory applied to this turgid hollow filament considered as an elastic thin rod [19]expresses the elastic flexural buckling load as a linear function of the bending stiffness EI[20], with *E* the Young modulus of the cell wall and *I* a geometrical parameter referred to as the second-moment of area. Knowing that the composition and structure of this protective shell change as a function of growth conditions[21] and environmental stress[22], the bending stiffness constitutes an interesting overall mechanical parameter to quantify the cell wall stress-induced responses.

Microfluidic devices specifically dedicated to the measurement of various mechanical parameters of filamentous walled cells such as the growth-stalling [15, 23] and protrusive[15, 24, 25, 26] forces have been developed during the last two decades[27] for various biological systems, i.e. *Schizosaccaromyces pombe*[23], the oomycete *Achlya bisexualis*[26], pollen tubes [24], and *C. albicans* hyphae [15, 25]. These passive methods rely on the deformation of elements of the microfluidic device by the forces produced by the growing cells. Measuring the bending stiffness required actuated devices. The seminal work of Sa-nati Nezhad et al. was dedicated to the deflection of single pollen tubes using hydrodynamic forces[28]. More advanced microfluidics tools employed “mother machine” derived devices, originally designed to study long-term growth and division patterns of bacteria, to measure the bending properties of arrays of filamentous *E. coli* and *B. subtilis* cells[29, 19].

We have elaborated on these previous works to build a device performing two complementary tasks: the positioning of *C. albicans* yeasts in a first (i.e. seeding) chamber and the application of hydrodynamic forces on the hyphae induced from these yeasts in a second (i.e. bending) chamber (figure 1a). An array of microchannels connects the two chambers and provides guidance for hyphal growth. Differently from a previous work[28], we implemented a tight mechanical maintenance of hyphae at the channel exit in order to maximize the contribution of the filament bending stiffness in the total observed deflection (i.e. to minimize the contribution of artifactual lateral displacement of hyphae inside the channels).

**Figure 1.**
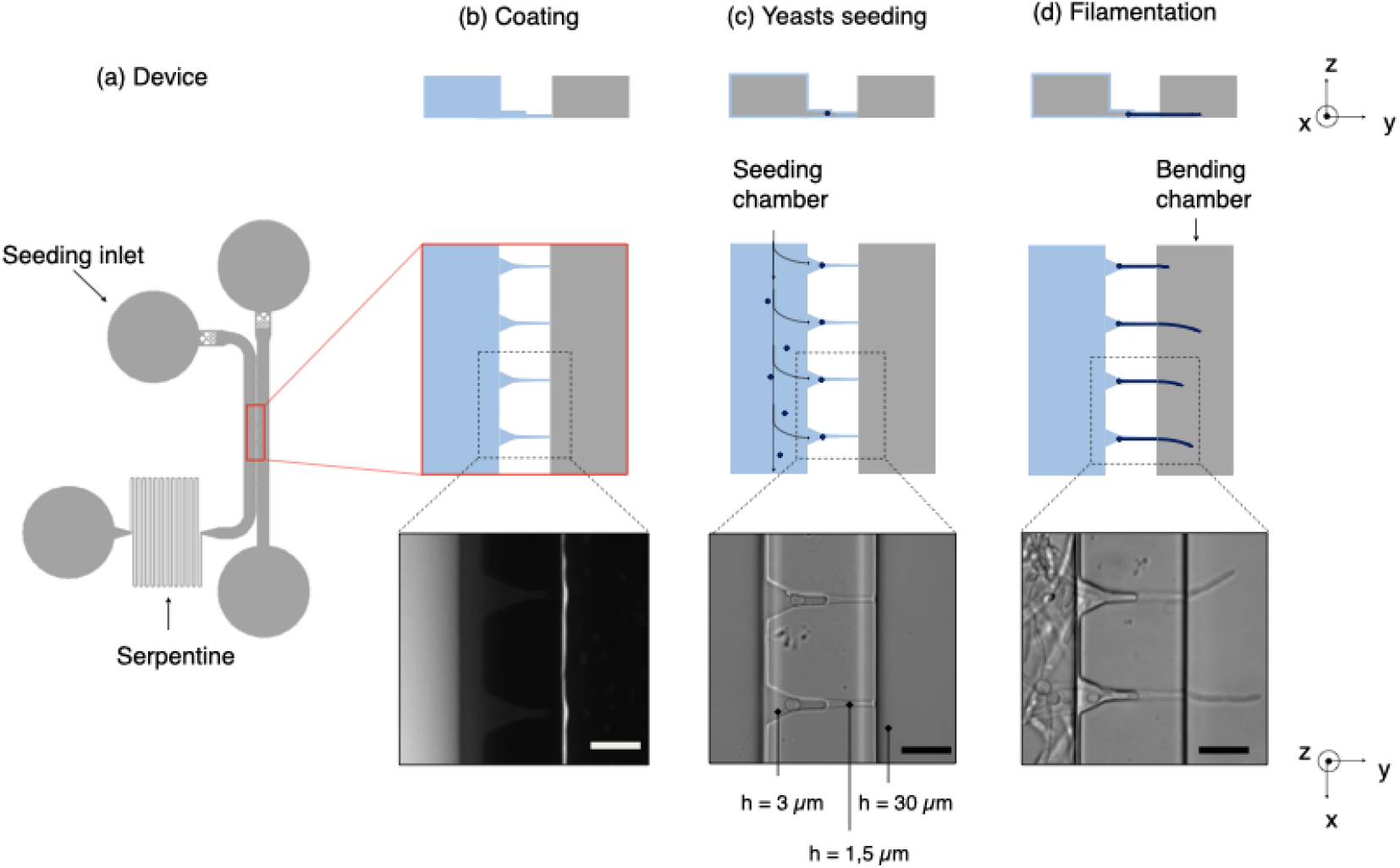
Design of the chip and experimental procedure. (a) Scheme of the microfluidic device containing 2 main chambers connected by microchannels, a leftward seeding chamber and a rightward bending chamber. A long serpentine is used to increase the hydrodynamic resistance of the seeding compartment and consequently to reach an efficient positioning of the yeasts in front of the microchannels. (b) Fibronectin coating (in blue on the schemes) into the seeding chamber. The bending chamber remains uncoated, as assessed from the localization of the PLL-FITC coating replacing non fluorescent FN in mock experiments (image, obtained from fluorescence imaging). (c) Trapping of yeasts at the entrance of the microchannels, followed by (d) hyphal growth in the bending chamber where they are subjected to shear flow. Scale bars: 20μm.

The deflection of hyphae was measured and analyzed through a theoretical framework based on the elastic beam theory. We carried out simulations based on finite element analysis to size the different parts of our device and to refine the mechanical modeling used to extract the bending stiffness from the deflection of hyphae.

We present here measurements of the bending stiffness of *C. albicans* hyphae in different conditions. We first assessed the action of caspofungin. This anti-fungal drug is known to target the synthesis of *β*-glucans, which predominantly occurs at new cell wall addition sites at the hyphal tip. Surprisingly, we observed dramatic reduction of the measured bending stiffness, suggesting that casponfungin affects the entire length of the hypha cell wall beyond the newly added material at the tip. The ability of the cell wall to remodel in response to an environmental stress could also be deduced from the observed decrease in bending stiffness following an hyperosmotic shock. Finally, a simple change in the cell wall thickness does not affect much its mechanical properties, as observed using a mutant of GPI-modified cell wall proteins. Overall, these findings provide new insights into how a hypha responds mechanically to various perturbations, and show that *C. albicans* is able to adjust its bending stiffness in a range of about one order of magnitude in response to environmental changes.

## Materials and methods

### Microfluidic chip fabrication

A first layer of SU-8 2002 (MicroChem) is spin-coated on a silicon wafer to obtain a final thickness of 1.5 μm. After a soft bake, this layer of negative photoresist is exposed to UV light thanks to direct laser writing lithography (Heidelberg, μPG101). Post-exposure bake and development with PGMEA are performed to obtain the first layer. The same process is repeated to add a second layer of 3 μm height. It is precisely aligned on the first one thanks to reference crosses on both layers. Then a third layer of SU-8 2035 is spread out on top of the mold in order to get the macro-chambers of 30 μm height. After an alignment step, the mold is then exposed to UV light through a chromium mask (MJB4 mask aligner). The heights of the mold were measured with a mechanical profilometer (Dektak).

The mold is first treated with trichloroperfluorooctylsilane (ABCR GmbH) deposited in vapor phase during 15 min in order to prevent adhesion of PDMS to the mold. Then PDMS is mixed with its curing agent (Sylgard 184 silicone elastomer kit) at the ratio 10:1 (w/w), degassed in bulk, poured on the mold, degassed again and finally cured at 75°C overnight.

The cured PDMS is then peeled off from the mold, inlets and outlets are punched with a 1.5 mm diameter biopsy punch (Kai Industries).

### Differential surface coating

After a step of surface activation using an oxygen plasma, PDMS chips are bonded on glass Petri dishes (Fluorodish, VWR). An adhesive coating only in the seeding chamber (left channel on the figure 1b) is then performed. For this purpose, a small drop of fibronectin bovine plasma (FN, Sigma) at 50 μg/mL is introduced in this chamber about 15 minutes after plasma treatment rather than immediately. This waiting time allows to tune the hydrophilicity of both chambers in order to favor spreading of the aqueous solution of FN in the seeding chamber while preventing it from invading the bending chamber through the microchannels. After 10 minutes, the chip is thoroughly washed with ethanol and dried.

### Yeast culture and hyphal induction

Most of the experiments have been performed using the SC5314 reference strain of *C. albicans*. A mutant with deletion of two GPI-anchored proteins Pga59 and Pga62 has also been used [30, 31]. Strains are stored at -80°C in YPD medium (1% yeast extract, 2% peptone, and 2% glucose in distilled water (w/v)) supplemented with 25% of glycerol (v/v). They are streaked on YPD agar plates (YPD medium supplemented with 1.5% of agar (15g/L of agar)) and kept for maximum 3 weeks at 4^°^C. Prior to each experiment, a preculture is done: a colony from the plate is put in 3mL of YPD liquid medium, and kept under agitation overnight at 30°C. Typically, 50μL of this solution diluted in 1mL of YPD corresponds to an optical density slightly below 1, corresponding to ≈10^7^ cells/mL. 5μL of the volume of this preculture (i.e. ≈10^6^ cells) is added to 495μL of the filamentation medium composed of 445μL of synthetic complete medium (i.e. SC medium, containing 0.67% yeast nitrogen base without amino-acids, 2% glucose, and 0.2% amino-acid mix[32] (w/v)) and 50μL of N-acetyl-D-glucosamine at 1 mg/mL. Overall, the dilution of the preculture is made so that about a thousand of cells are inserted in the seeding chamber.

### Bending hyphae under tangential flow

Hyphal growth takes place into an incubator regulated at 37°C until most hyphae have crossed the microchannels and have started to invade the bending chamber. Usually a few hours are necessary to grow filaments that protrude 30μm approximately. The chip is then ready for bending experiments, which take place at room temperature using purified water (Millipore Milli-Q lab water system) to impose the tangential flow. In that aim, the chip is connected to a pressure-based flow controller (Flow EZ, Fluigent) and a flow sensor (Flow Unit, Fluigent) is added upstream. Water flow rate can then be controlled in the bending chamber, and flow rates ranging from 0 to 40μL/min (equivalent in our system to pressures ranging from 0 to 80mbar) are incrementally used to deflect the filaments.

The above protocol is used to measure hyphae deflexion in control situations. To assess the effect of caspofungin, it is followed by the replacement of the medium present in the whole chip by a fresh SC medium containing the desired concentration of the drug (in our case 1μg/mL, prepared from 0.5μL of a stock solution at 1mg/mL mixed with 495μL of the filamentation medium). The device is then put again 30 min into the incubator before the same protocol as for control experiments is applied to measure the hyphae deflexion.

Regarding sorbitol, no incubation step is required and the concentration in sorbitol of the water bending solution, namely 2.5M (455g/L of sorbitol powder) is set the same than in the seeding chamber to provide uniform osmotic conditions all over the device.

### Numerical simulations

3D simulations were performed with Comsol Multiphysics. First, simulations of the flow to get the velocity profile in the bending chamber was carried out. The flow calculated in a simplest model of the bending chamber, i.e. a rectangular channel, was compared to calculations using a more realistic design which includes the microchannels (ESI† Fig.S1a-c). A variation in the velocity profile of less than 5% was found, thus showing the negligeable contribution of microchannels in the flow distribution near the surface of interest. We have thus considered in further simulations the flow in a rectangular channel without taking into account the rest of the device.

Then rigid cylinders were introduced in our simulations in order to model the bending of filaments under the flow. We chose to implement rigid cylinders instead of tubes for the sake of computation time, the thin meshing required for a tube being useless for this much larger spatial scale computation. We used the fluid-structure interaction module, considering that one end of the cylinders was fixed to the wall to reproduce the immobilization of hyphae by the microchannels.

## Design of the microfluidic chip

The microfluidic device is composed of 2 main compartments, i.e. the seeding and bending chambers, separated by an array of microchannels (figure 1a).

The first step of this experiment consists in positioning the yeasts in front of the microchannels prior to their filamentation (figure 1c). High-efficiency trapping of cells was previously obtained with designs of various complexity[33, 34, 35]. Here we chose to introduce a long serpentine at the exit of the seeding chamber. Its length and width were chosen so that its hydraulic resistance is equivalent to that of microchannels. This design allows a large fraction of the flow to be diverted toward the bending chamber through the microchannels, driving about 3/4 of the yeasts in that direction where they will get trapped into small pockets designed to accommodate single yeasts.

The microchannels were sized smaller than the characteristic size of yeasts (5μm), so that these cells will not be able to penetrate further into these channels. Once most of the traps are occupied, the resistance on this path will increase due to yeasts blocking the channels. Any excess of cells will be thus redirected towards the outlet.

Beyond their filtering effect toward yeasts, the role of microchannels is threefold. First, they provide initial directional guidance, leading hyphae to emerge as perpendicularly as possible to the edge of the bending chamber (figure 1d). Second, they prevent these filaments to move laterally under the flow: it is essential to avoid this contribution by adjusting as much as possible the dimensions of the microchannels to the size of hyphae. Third, they localize the exit of hyphae on the bottom surface of the bending chamber. We observed that about one third of the filaments continue to growth close to this surface. We thus choose, contrarily to [28], to work within a single focal plane of observation, which simplify both data acquisition and analysis procedures and increases significantly the throughput of a single experiment.

To fulfill these aforementioned functions at the best, we designed funnel-like channels characterized by 2 successive heights of 3μm and 1.5μm (ESI† Fig. S2). We observed that a 3μm height is well adapted to induction and guidance of the germ tube arising from the trapped yeast. However the space around the filament was too large to prevent its lateral or vertical movements. A reduction of the height in the final portion of the microchannel suppressed these unwanted effects.

Besides, backward movements had to be prevented. In the case of pollen tubes, this was performed by introducing kinks in the growth channel[28]. In the present study, FN coating of the seeding chamber and the use of narrow channels were enough to immobilize both the mother yeast cells and the hyphae against backward displacements under flow.

Lastly, while hyphae need to be fixed on one end, bending experiments also require them to be able to move and deflect under the flow in the bending chamber. It is thus crucial to avoid adhesion between hyphae and the substrate in these experiments. This is the reason why the FN coating was limited to the seeding chamber and therefore the PDMS was left uncoated in the bending chamber (see methods).

The length of these microchannels was set to 40μm to ensure a guided filamentation in a relatively short time, considering an elongation rate of the hyphae around 0.3μm/min. In order to maximize the number of filaments inside the bending chamber while avoiding a crosstalk in the velocity profiles from one hypha to its nearest neighbor, we determined the deflection of a single filament and compared it to the value obtained when the filament was surrounded by several others separated by various distances (ESI† Fig. S3a-b). We finally obtained an optimized distance of 40μm between microchannels.

## Theoretical model: elastic deformations of a beam

As in previous works [19, 28, 29], we used the hollow tube deflexion model to obtain the bending stiffness of hyphae. We also implemented some refinements of this general modeling frame to better describe our system. We detail below the overall approach that we have set-up to retrieve the mechanical properties of hyphae from their deflection under flow.

### Filaments

We considered hyphae as hollow cylinders with a radius *r* and a cell wall thickness *t_W_*. The second moment of area *I* (in m^4^) with respect to the flow direction axis is expressed as follow for these filaments : *I* = *πr*^3^*t_W_*.

### Force exerted on the filaments

The flow is laminar and can be precisely computed in the rectangular bending chamber. As already stated above, only the filaments staying close to the surface are taken into account. In this frame, the force per unit length *f* exerted by the flow on a cylinder lying on a surface [36, 19] is expressed as :

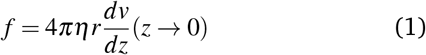

where *η* is the viscosity of the liquid (Pa.s), r is the radius of the filament (m) and *dv/dz* is the gradient of velocity along the z direction close to the bottom of the channel (s^-1^).

Indeed, in this configuration, numerical simulations show that, close to the surface (where z goes to 0), the gradient of velocity along the y direction, i.e. *dv/dy*, is negligible compared to *dv/dz*. However, *dv/dz* is not constant along the *y* direction (see the spatial profiles of both *dv/dy* and *dv/dz* along the *y* direction in ESI† Fig.S4), meaning that the force *f* is not constant along *y*. This spatial variation of *f*(*y*) was taken into account to compute *EI* from the hydrodynamic force and the deflection. This point is detailed in the next section.

### Expression of the bending stiffness

Assuming small elastic deformations and a uniform force per unit length *f* on the filament along the *y* direction (figure 2a), the Euler-Bernouilli beam theory for a cantilever beam subjected to a uniformly distributed load leads to the following relationship between the beam’s maximal deflection *δ*, the applied force *f* and the bending stiffness *EI* [37, 19]:

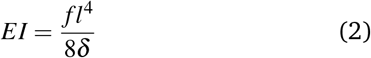

with:

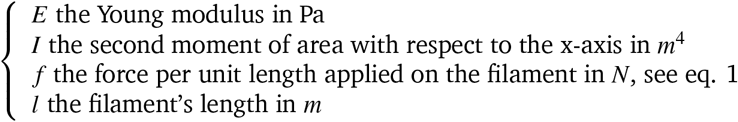

**Figure 2.**
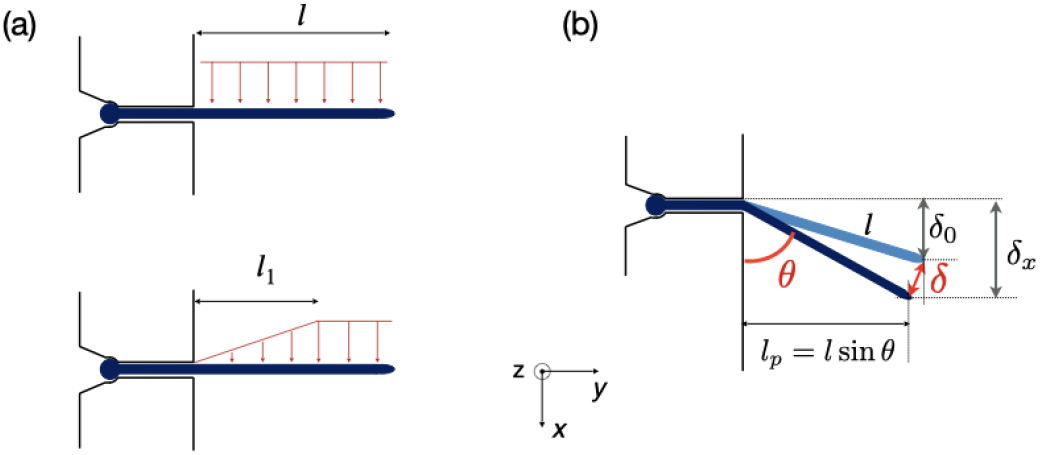
Forces exerted on a single hypha and deflection. (a) Scheme of hydrodynamic forces exerted on a hypha. Model considered in [19] (top) versus model developed in this study, taking into account the variation along *y* of the hydrodynamic forces (bottom). The length l_1_ is deduced from Comsol simulations, see ESI† Fig.S4. (b) Geometrical parameters characterizing a hypha of length *l* tilted by *θ* under the action of hydrodynamic forces.

Although not considered in [28, 19, 29], and as mentioned in the previous section, the force *f* is not constant along the length of the hyphae (i.e. along the *y* direction, see ESI† Fig.S4). The best linear approximation of these simulations is to consider the force not as a constant but as an increasing load over a length *l*_1_ before reaching a plateau for *l* > *l*_1_ (figure 2a). The value for l_1_, determined using Comsol, is around 15 μm. Considering that the mean value of the hyphal protruding length considered in this study is close to 30μm, this refinement of the model is particularly pertinent as it would account for an overestimation of *EI* by 10% for this length, and by about 22% for a length of 20μm. Using this refinement of the model, equation 2 becomes

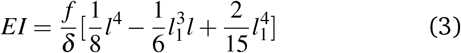

for filament lengths *l* > *l*_1_.

### Force-flow relationship

As shown in eq.2 the bending stiffness *EI* is deduced from the *δ*(*f*) variation. Experimentally, we record instead the *δ*(*Q*) curves and then deduce *f* from the measure of *Q*.

The relation between *f* and *Q* can be obtained from the Poiseuille velocity distribution for a liquid flowing in a channel of width *w* and height *h*: 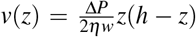. From this *v*(*z*) relationship, we obtain 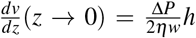. On the other hand, 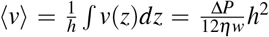, which leads to 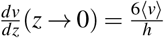. From the straightforward relation 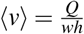, we finally obtain

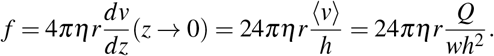

The overall angular tilt induced by the applied hydrodynamic force, eventually superimposed to an initial growth tilt, implies different geometrical corrections to deduce the deflection value *δ* (*Q*) (figure 2b). First, an initial deflection *δ*_0_ may result from the eventual existence of a flow-free initial tilt, which should be subtracted to the overall deflection *δ_x_* measured along the *x* axis. Next, *δ*(*Q*) = (*δ_x_* – *δ*_0_)/sin *θ*, with *θ* the angular tilt induced by the flow. Finally, the hydrodynamic force is exerted only over a length *l_p_* = *l* sin *θ*, meaning that a tilted hypha will be submitted to a force reduced by sin *θ*. Considering equation 3 and the above corrections, the general expression of *EI* is finally:

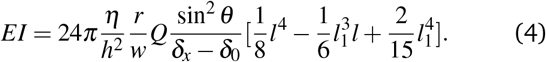

In practice, the term 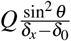 will be deduced for each hypha initially deflected by *δ*_0_ from the best linear fit of the variation of the deflection *δ_x_* and the associated angular tilt values *θ* for a series of increasing flow rates *Q*.

## Results

### Measurement of the bending stiffness of hyphae

We applied increasing and decreasing flow rates on hyphae in the bending chamber. The time allowed to each step was set to about 15 seconds, which is a good trade-off between reaching a stable flow and limiting the duration of the whole experiment, in particular to avoid any drift in hyphal length. The maximum flow rate was chosen to remain in the limit of reversible deformations, which was assessed by performing a full cycle of flow increase and decrease and by checking the absence of major hysteresis. We computed then the bending stiffness *EI* by taking into account the points obtained during the increasing part of the flow cycle.

Typical images and data resulting from such experiments are shown in figure 3a-c. In this particular example, the hypha was initially tilted by *δ*_0_ (figure 3a). Figure 3b shows the raw deflection values *δ_x_* after a full cycle of flow increase and decrease (top graph) and the linear fit of the points leading to term *δ* values for the raising part of the cycle only. The calculation of the bending stiffness *EI* uses the inverse of the slope displayed in this later figure, i.e. *Q*/*δ* (see equation 4).

**Figure 3.**
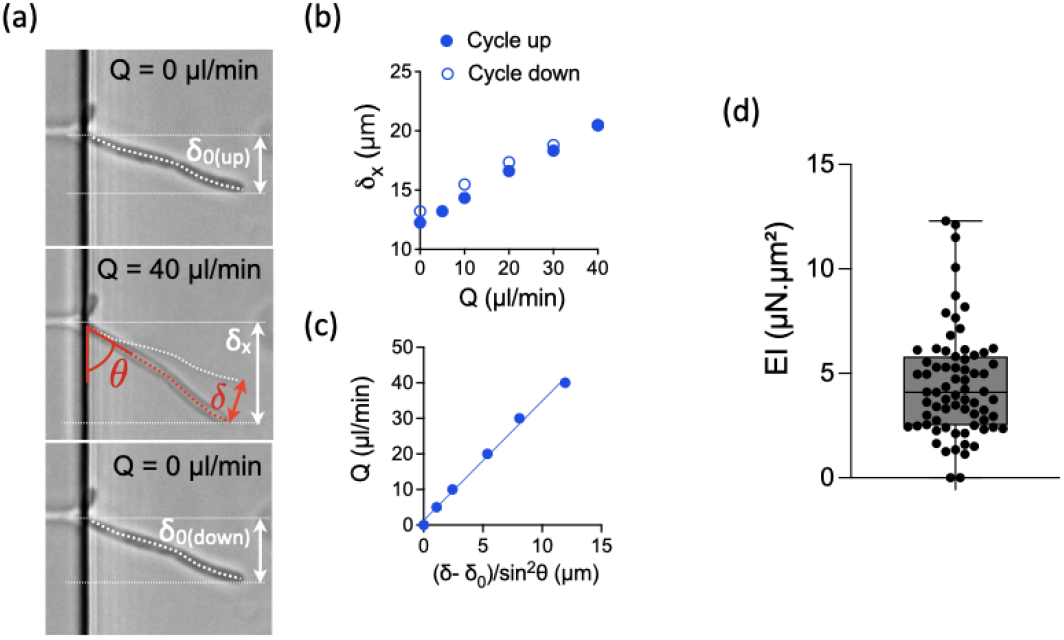
Bending properties of hyphae from the reference strain SC5314. (a) Images of an hypha initially deflected by *δ*_0_ before (top), during (middle) and after (bottom) flow application. The initial tilted position of the hypha is indicated in the middle image. The amplitudes of the deflection *δ*_0_, *δ_x_* and *δ* are indicated according to the nomenclature used in figure 2. (b) Example of deflexion curves obtained for this hypha. Top: deflexion *δ*_*x*_ obtained during increasing (solid circles) and decreasing (open circles) flow rates Q. Bottom: deflection data formatted to show the linear fit from which the bending stiffness EI will be deduced for each hypha (see equation 4). Only the data corresponding to increasing flow rate Q are considered. (c) Distribution (mean±SD = 4.5±2.5*μN.μm*^2^) of bending stiffness EI deduced from the slope of the linear variation shown in (b). n=69 hyphae.

For the reference strain SC5314 of *C. albicans*, we measured a mean bending stiffness *EI* = 4.5 ± 2.5*μN.μm*^2^ (n=69, figure 3c). This value is about 2 orders of magnitude above the bending stiffness reported for *E. coli*[19, 29, 38]. Considering the measured diameter values for each hypha and the assumption of a cell wall thickness of 230 nm[30], we estimated the longitudinal Young modulus of the reference strain SC5314 to be 6.4±3.4 MPa, interestingly in the same range as the values obtained for *E. coli* [39]. The literature provides a few values of this parameter for different yeast species. The fission yeast displays a Young modulus of ≈30 MPa[40, 23, 41]. For *S. cerevisiae*, the reported values are distributed in a wide range, from around the MPa[42, 43, 44] to hundreds of MPa[45, 46]. Values of the Young modulus of *C. albicans* are also quite distributed between very low (i.e. ≈100-200kPa) [47, 48, 49] to values in the MPa range [50, 51, 52]. The values we obtain with our method are thus coherent with the order of magnitude reported in part of the literature, whose inconsistencies remain to be elucidated.

### Influence of caspofungin treatment on bending stiffness

Caspofungin belongs to a new class of antifungal agents, the echinocandins. This antifungal drug is known to inhibit the synthesis of *β*-1,3-glucan, which is a main component of the fungal cell wall. This synthesis is particularly active at the apex. It is thus expected that the cell wall at the tip should be the major target of caspofungin. However, this antifungal agent seems to also affect the cell wall outside the location where new synthesis occurs, i.e. the mature cell wall[53]. Considering that any disturbance of the cell wall should affect the bending stiffness of hyphae, we decided to explore the effect of this antifungal agent in our system.

Filaments were first subjected to a cycle of increasing and decreasing flow rates to obtain a reference stiffness for each filament in a medium devoid of caspofungin. Then, as described in the Materials and Methods, the measured hyphae were incubated for 30 minutes at 37°C in a culture medium supplemented with caspofungin at a concentration of 1μg/mL. Such an acute drug application at a concentration well above the minimum inhibitory concentration (MIC) of 0.125μg/mL in glucose-rich medium [53] was chosen to maximize the chance to see an effect of caspofungin while limiting changes in hyphal length and overall morphology (i.e. branching) and to prevent any adaptation mechanisms through the activation of cell wall compensatory pathways [54]. Control experiments (see ESI† Fig. S5) without caspofungin in the bending medium were performed, showing that the 30 min incubation and medium change steps does not modify the bending stiffness of filaments. The behavior of most filaments could be followed before (n=18) and after (n=21) the application of the drug (see an example in figure 4a). For a minority of filaments however (n=5), we got data for only a single condition, the one related to caspofungin being more represented (n=4). In this case, hyphae previously immobile despite the applied flow were released from their adhesion to the surface after the introduction of the antifungal agent. We chose in figure 4b to present the distribution of all data together with the ratio of paired data. The distribution of paired data can be found in the supplementary figures (see ESI† Fig. S6a).

**Figure 4.**
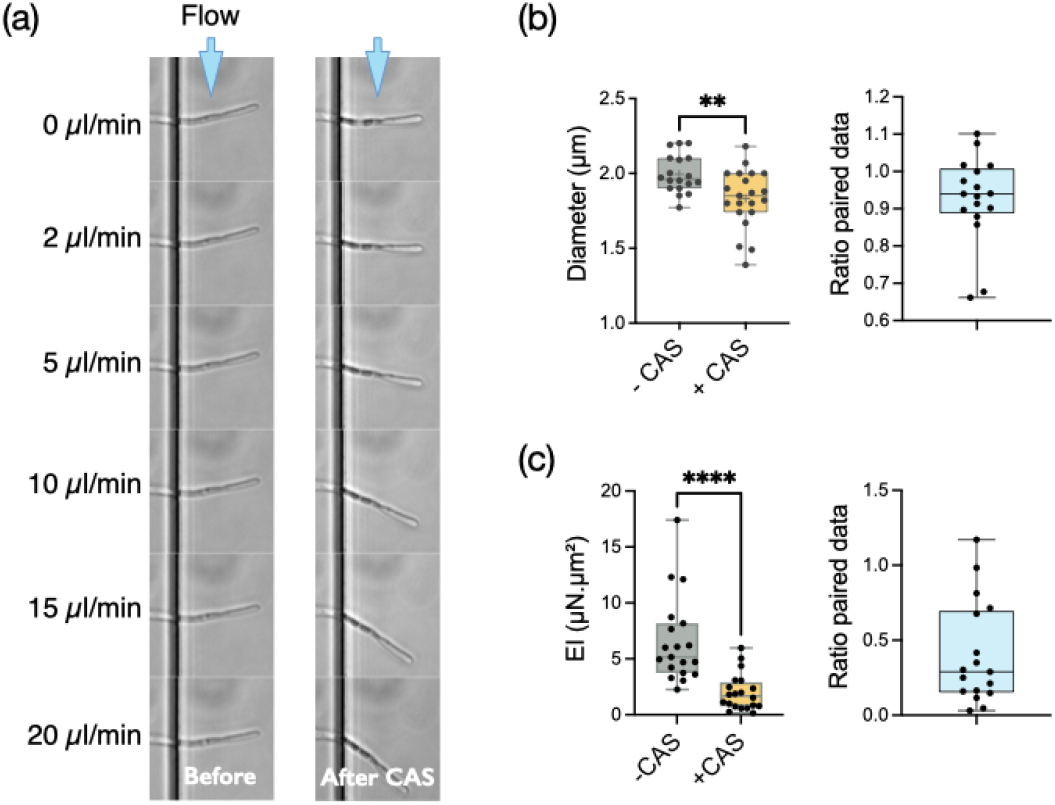
Consequences of an antifungal treatment using caspofungin at 1μg/mL for 30min on the reference strain SC5314. Graphs display means and standard deviations. (a) Images of a hypha bending under flow before and after caspofungin treatment. (b) Diameters. Left: distribution of diameters before (1.99±0.13μm, n=18) and after (1.83±0.20μm, n=21) antifungal treatment; p=0.0081 (Mann-Whitney test); Right: diameter ratio for paired data (0.93±0.12, n=17) (c) Bending stiffness *EI*. Left: distribution of the bending stiffness *EI* before (6.54±3.84μN.μm^2^, n=18) and after (1.98±1.63μN.μm^2^, n=21) antifungal treatment, p<0.0001 (Mann-Whitney test); Right: distribution of *EI* ratio for paired data (0.117±0.0283, n=17).

We first observed that this concentration of caspofungin reduces slightly the diameter of hyphae, from 1.99±0.13*μ*m without caspofungin to 1.83±0.20*μ*m with this drug (figure 4b). of the hyphal tip. But the main effect of caspofungin application is a reduction by more than a factor of two (ratio 0.40±0.34) of the bending stiffness *EI* of hyphae (figure 4b). The mean reduction of the diameter *d* by a factor of 0.926 induced by caspofungin would lead to a reduction of the second moment of inertia *I* ∝ *d*^3^ of ≈ 0.79. The caspofungin-induced morphological effect can thus only partially account for the bending stiffness variation. We thus conclude that caspofungin affects the mechanical properties of the cell wall at the hypha scale.

These filaments being composed of mononucleated compartments separated by septa, we wanted to check if the presence of these internal cross-walls could be a preferential target for the action of caspofungin. The length of individual compartments is typically 20μm while the hyphae protrude from 10 to 55μm in the bending chamber. First we did not observed angled hyphal shapes after caspofungin application that would suggest the presence of bending points at the septa. Then we looked for a variation in the bending stiffness as a function of length, as any specific impairment of the septa would lead to an additional increase of the deflection. We did not measure a significant variation of EI as a function of hyphal length (ESI† Fig. S8). We thus concluded that a large part (i.e. ≈2/3) of the reduction of the bending stiffness after caspofungin application should be attributed to a reduction of the cell wall Young modulus E, or the cell wall thickness, or most probably a combination of both.

### Influence of sorbitol treatment on bending stiffness

In *C. albicans* yeast, osmotic shock induces a volume decrease within seconds, associated with major changes in cell wall architecture. These structural changes occur prior the activation of high-osmolarity glycerol (Hog1) or cell integrity signaling pathways which takes tens of minutes to manifest themselves[55]. This shrinkage of the cell wall perimeter in yeast, and its reversibility when returning to an osmoneutral medium, shows its highly elastic properties, attributed mainly to the spring-like properties of the *β* 1,3-glucan chains found in the inner cell wall[56]. The time scale of these changes being too fast to allow a modification of the cell wall overall volume, the almost instantaneous reduction in cell diameter leads at very short term to an increase in the cell wall thickness, and specifically the inner *β*-glucan and chitin layers[55].

However, if the cell wall properties of yeasts under osmotic stress have been explored, there is still little reports on hyphal properties under such stress. We thus set up experiments to measure the bending stiffness of hyphae submitted to an hyperosmotic shock provided by 2.5M of sorbitol, a polyol compound known to increase the osmolarity of aqueous solutions[57]. Hydrodynamic forces were applied within about 5 minutes after sorbitol medium replacement.

As expected, the hypha diameter decreased significantly (i.e. by a reduction factor 1.28) under the application of sorbitol (figure 5a). Similarly to what we observed using caspofungin, the reduction of diameter tends to free previously adherent hyphae, leading to more data for the condition with sorbitol (see ESI† Fig. S6b for the distribution of paired data only). The analysis of hypha deflection revealed an even more dramatic decrease of the bending stiffness by a factor of about 7 (figure 5b). The extent of this change cannot be attributed only to the hypha’s diameter reduction which alone would lead to a factor of only 2 (i.e. 1.28^3^) in the reduction of *EI* through the r^3^ dependence of the second moment of area *I*. Moreover, the contribution of *I* = *πr^3^t_W_* in the measured *EI* values might be even reduced if taking into account an increase of the cell wall thickness t_W_ proportional to hyphal diameter reduction (considering the reasonable hypothesis of a conservation of the biomass of the cell wall). We conclude that most of the changes in the bending stiffness *EI* should therefore be attributed to a modification of the cell wall mechanical properties induced by a rapid remodeling of the cell wall.

**Figure 5.**
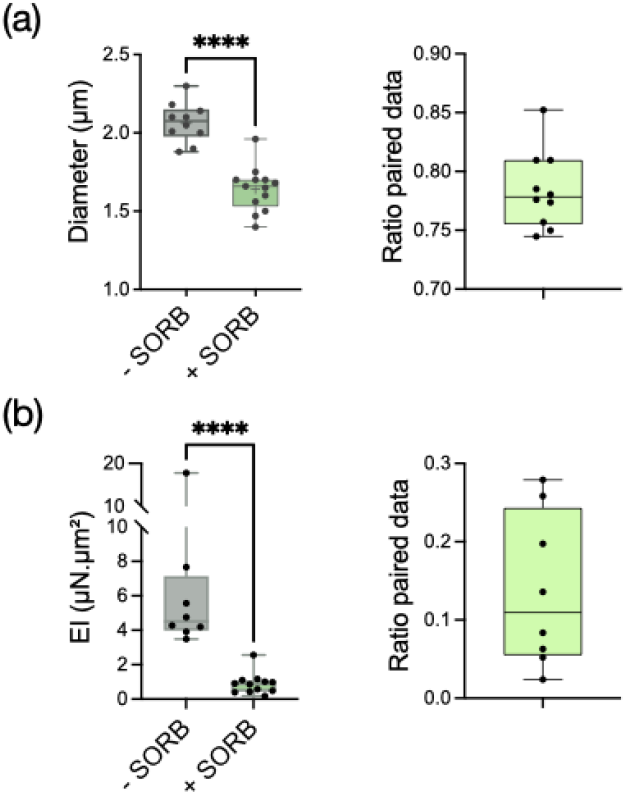
Behavior of the reference strain SC5314 before and after an hyperosmotic shock provided by 2.5M of sorbitol. Graphs display means and standard deviations. (a) Left: distribution of diameters before (2.07±0.13μm, n=10) and after (0.142±0.039μm, n=13) sorbitol application, p<0.0001 (Mann-Whitney test); Right: diameter ratio for paired data (0.78±0.033, n=10). (b) Left: distribution of the bending stiffness *EI* before (6.46±4.77μN.μm^2^, n=8) and after (0.888±0.615μN.μm^2^, n=12) sorbitol application, p<0.0001 (Mann-Whitney test); Right: distribution of EI ratio for paired data (0.14±0.099, n=8).

### Study of a cell wall mutant

Most proteins in the yeast cell wall are glycosylphosphatidylinositol-anchored proteins (GPI-modified proteins) which are covalently linked to it. Simultaneous deletion of the genes for two of these cell wall proteins, Pga59 and Pga62, does not have a major impact on hyphal growth. However, it induces a modification of the cell wall thickness [30]. More precisely, the total thickness measured by transmission electron microscopy increases from 230 nm in the SC5314 wild type strain to 290 nm for the Δ*pga*59Δ*pga*62 mutants, while the inner wall thickness evolves from 150 to 230nm.

We tested if these modifications would affect the bending stiffness of this mutant (Figure 6). Although non significant, the increase in the mean value of the bending stiffness measured for the Δ*pga*59Δ*pga*62 mutant strain as compared to the reference strain is similar to the ratio between the thickness of their total cell wall (1.2 compared to 1.26, respectively). We conclude that when the diameter of the hypha is not modified, as it is the case here, a change in the cell wall thickness affects the mechanical property, but the linearity between this morphological parameter and *EI* prevents any major change in the mechanical properties of the cell wall.

**Figure 6.**
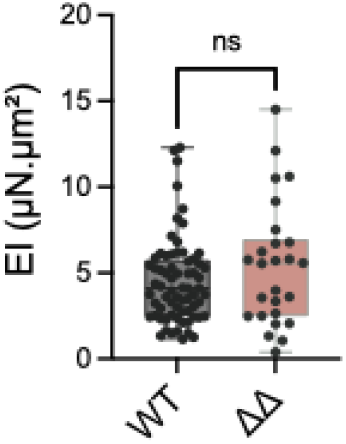
Distribution of the bending stiffness *EI* of the reference strain SC5314 (WT, 4.50±2.49μN.μm^2^, n=69) and of the Δ*pga*59Δ*pga*62 mutant (ΔΔ, 5.44±3.60μN.μm^2^, n=26), p=0.329 (Mann-Whitney test).

## Discussion

We have developed a microfluidic device capable of measuring the bending strength of individual *C. albicans* hyphae by applying hydrodynamic forces. The analysis of bending data implied some experimental precautions, that we have implemented (e.g. maintaining the base of the hypha during flow application). It also led to the adaptation of a general model by taking into account the geometry of our system (edge effects for the calculation of the gradient of the hydrodynamic force) as well as the deviations observed with respect to a canonical situation (initial inclination of the hypha, lateral displacement at the exit of the microchannel).

The measured bending stiffness is the product of an elastic modulus and of a term combining various morphological characteristics of the hyphae (length, diameter, wall thickness). It is tempting, and we have given some examples earlier in the text, to compare these values to those obtained from AFM indentation measurements. However, several remarks must be made at this stage. First of all, the elastic modulus obtained by indentation combines the moduli associated to the 3 directions of space. Milani et al. have showed for the cell wall of living shoot apical meristems that indentation experiments were essentially sensitive to the normal modulus (i.e. the module along the direction of indentation), which contributes 4 times more than the longitudinal and transverse moduli to the global apparent modulus [58]. In contrast, the wall is tangentially stretched during bending deformations. In other terms, the tangential modulus should be a major component of the apparent elastic modulus involved in the bending stiffness. Moreover, considering the fibrous structure of the cell wall (in both fungi and plants), this tangential modulus should be higher than the two others. A normal force will indeed essentially deform the soft matrix surrounding the fibers while the stiffness of the fibers, higher than their surrounding matrix, will mostly be responsible for the strain induced by a tangential stretch. We therefore expect to deduce from our bending measurements higher elastic moduli than those measured by AFM.

Note that AFM measurements are very sensitive to the amplitude of indentation. It must remain low at the risk of probing the turgor pressure together with the cell wall rigidity. This is besides an elegant way to measure these two properties in a single experiment[46]. However, this point is rarely explicitly mentioned or discussed in the literature. The two modalities of the AFM technique (normal direction and amplitude of the indentation) may have opposite effects on the estimation of the Young modulus: one would tend to minimize the resulting effective modulus (by giving an higher weight to the lowest, i.e. normal modulus), and the other would on the contrary lead to overestimate it (from a possible contribution of the turgor pressure in the case of a too large indentation depth). It is thus difficult to compare with precision our measurements with those published in the literature. However, despite a very wide disparity in the moduli published for *C. albicans*, we find that the values deduced from our measurements are in the high range, consistent with the above discussion.

In this study, we used a concentration of caspofungin well above the minimum inhibitory concentration (MIC) and an incubation time of 30 min, conditions that we identified in control experiments as capable of altering the adhesive properties of the yeast form of *C. albicans* while not affecting cell viability (not shown). This concentration is also expected to be lower than the concentrations giving rise to the “paradoxical effect” [59] characterized by activation of chitin synthesis in response to inhibition of *β*-1,3-glucan but also *β*-1,6-glucan[60] production. The deflection measurements show different effects related to caspofungin. The first one relates to a loss of hyphal adhesion, as evidenced by the detachment of filaments previously anchored on the PDMS surface during control experiments (i.e. before casponfungin application), consistent with our observations on the yeast form. This property of caspofungin has been previously reported from AFM adhesion strength measurements on yeast [61] and hyphae [47], at concentrations similar and up to 3 orders of magnitude lower than those used in the present study. The other effect is a slight decrease in hyphal diameter after application of caspofungin, a property that could contribute to the observed loss of adhesion, in complement to the hypothesis of a lowering of adhesin expression in response to caspofungin[47].

The major effect of caspofungin observed in the present study remains the drastic decrease in the bending stiffness. This can come from two simultaneous effects: the decrease of the wall thickness (“I” component of the bending modulus) and a weakening of the Young’s modulus of the wall. Wall thinning has been suggested in [62] and [63] where it is stated that strains with increased caspofungin susceptibility have thinner walls. This hypothesis was recently verified experimentally by direct measurements for a concentration of 5μg/mL and 30 minutes of incubation[64]. This phenomenon is accompanied by cell bulging, more and more pronounced at long times (> hour), which we also sometimes distinguish at the hyphal tip (see figure 4). The observed decrease of the Young’s modulus of the wall is consistent with the concentration used, too low to induce a synthesis of chitin aiming at counteracting a weakening of the cell wall, but strong enough to trigger a loss of viability linked to cell wall or plasma membrane damages[53]. Our results are also in agreement with the results of AFM measurements performed for concentrations similar to ours[61]. Concentrations 20 to 2000 times lower over longer times, however, seem to give opposite results[48, 47]. This decrease in Young’s modulus at the scale of a hypha may however be surprising, as the inhibition of *β*-glucan synthesis was expected to be localized only at the tip. However, our results are in line with the work of Hao et al.[53] who observed in *C. albicans* that dividing yeasts exposed to caspofungin mother and daughter cells exhibited similar vitality loss, as evidenced by methylene blue staining intensity. These authors suggested a continuous cell wall remodeling process as an explanation to these results.

The significant decrease in hyphal diameter subjected to 2.5M sorbitol reflects the expected loss of *C. albicans* turgor pressure following an hyper-osmotic shock. Turgor pressure should not be involved in the calculation of bending stiffness at low strains. This assumption is based on finite element simulations on one hand[28], and on theoretical considerations based on the main argument that a bending deformation of a pressurized tube does not modify its internal volume[65, 66]. However, we prefer to leave open the question of a possible contribution of the turgor pressure for future works, in particular because the deflections observed under sorbitol application might not fall anymore into the low deformation range considered theoretically in [66]. Besides, the diameter reduction is expected to trigger a reconfiguration of the wall structure, which could even resemble to a destructuring considering the abruptness of this event. This impairment of the cell wall structure could contribute to the decrease of the Young’s modulus value we observed.

## Conclusion

Many questions in plant and fungal biology address the mechanics of pressurized filaments. Complementary to local methods such as AFM, the strategy developed here allows an access to the mechanics of *Candida albicans* at the hypha scale through the measurement of their bending stiffness. This method is generic and could be applied to the hyphae of multiple species of filamentous fungi by a simple tuning of the dimensions of the microfluidic device. In this first study dedicated to *C. albicans*, we were able to highlight the consequences in hypha mechanical properties of an anti-fungal drug widely used in clinic and of an hyperosmotic shock, while we noticed that modifications of the wall structure do not always lead to significant changes. Our work thus shows the complementarity of pharmacological, structural and biomechanical approaches in the study of the biology of the cell wall of hyphae.

## Conflicts of interest

There are no conflicts to declare.

## Acknowledgements

We warmly thank Arezki Boudaoud for his careful critical reading of the manuscript. This work benefited from the technical contribution of the joint service unit CNRS UAR 3750. The authors would like to thank the engineers of this unit for their advice during the development of the experiments. This work was supported by the European Research Council Advanced Grant 321107 “CellO” and the Institut Pierre-Gilles de Gennes (“Investissements d’avenir” program ANR-10-IDEX-0001-02 PSL and ANR-10-LABX-31, and ANR-10-EQPX-34).

## Notes

### Competing Interest Statement

The authors have declared no competing interest.

