## Supplementary data for "Bending stiffness of *Candida albicans* hyphae reflects adaptive behavior of the fungal cell wall"

### 1 - Numerical simulations to validate the theoretical model

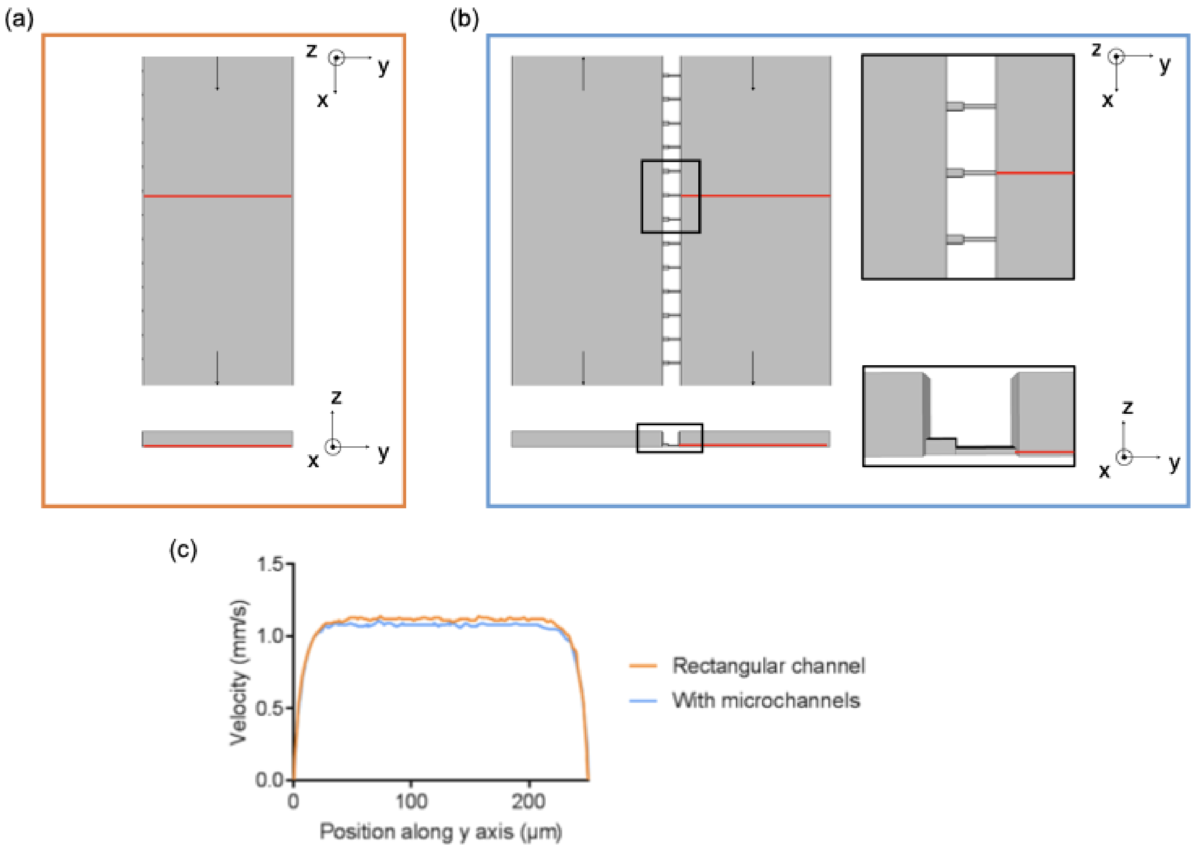


Figure S1 - Numerical simulations with Comsol to study the flow in the bending chamber. Comparison of the flow in a simple rectangular channel (a) and in the bending chamber included in the whole chip, taking into account the presence of microchannels (b). (c) The velocity (mm/s) is plotted along the red line in both situations as a function of y position (μm) close to the hyphae position at a fixed z position ($z = 1$ μm).

### 2 - Two height microchannels

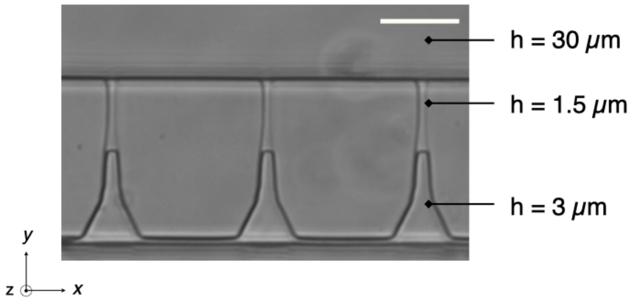


Figure S2 - Phase contrast image of microchannels. The height of each portion is indicated, as well as the height of the bending chamber. Scale bar: 20μm.

#### 3 - Influence of the distance between microchannels (filaments) on the measured deflexion under flow

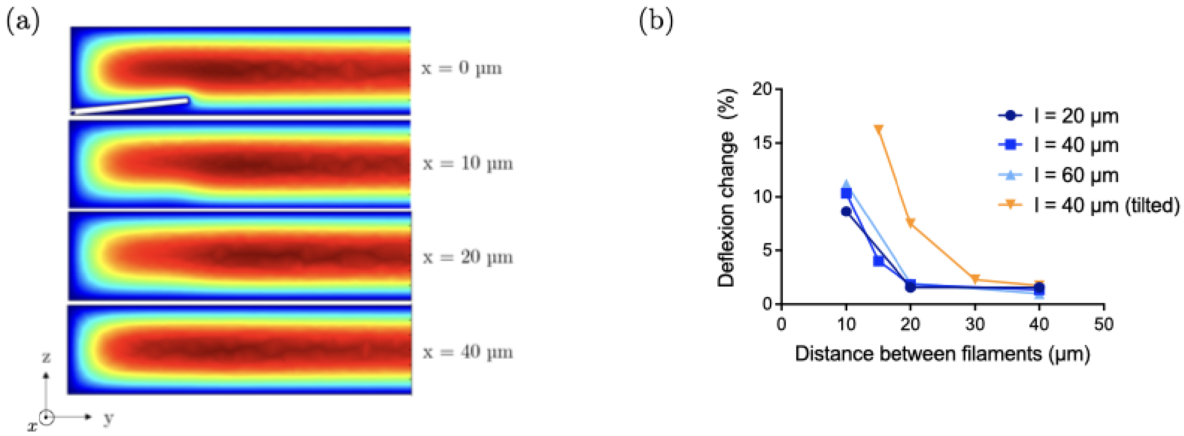


Figure S3 - Optimization of the distance between microchannels. (a) Velocity profiles in the bending chamber at different $x$ positions from a 40µm long tilted cylinder along $z$ modeling an hypha non adhering to the surface. This situation is associated to the maximal flow perturbation (see (b)). The changes in the velocity profile observed close to the filament vanished at a distance of 40µm. (b) Difference in deflection values between a situation where several filaments of length $l$ are spaced by a distance d along the $x$ axis compared to a situation with a single filament. The case of a tilted filament compared to the glass slide is also shown.

#### 4 - Numerical simulations of velocity profiles

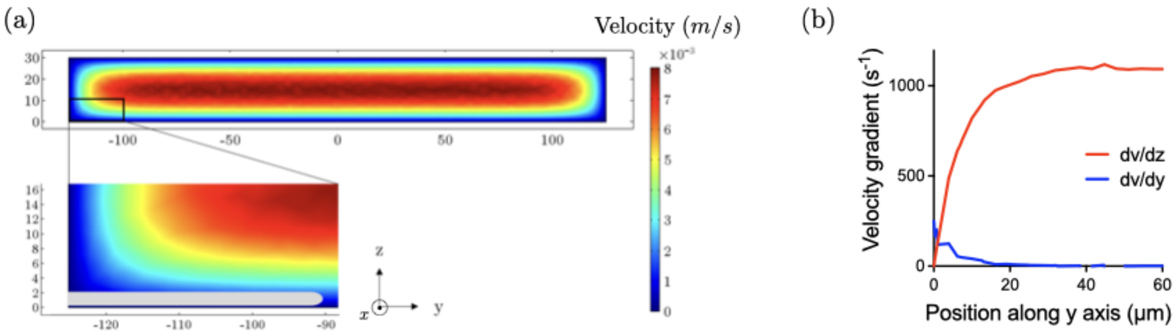


Figure S4 - Flow in the bending chamber: numerical simulations with Comsol. (a) Velocity profile in a rectangular channel of width L = 250 µm and height h = 30 µm for an inlet velocity of 5 mm/sec. The insert shows an enlargement of the area of interest, where the hyphae come out of the microchannel (in light grey). (b) Velocity gradient (in $\sec^{-1}$) $dv/dz$ and $dv/dy$ along the y axis at a height z = 1µm. For filaments whose length range from 20 to 50 µm, gradient along y is negligible compared to the one along z. Moreover, it appears clearly that $dv/dz$ varies as a function of $y$ close to $y = 0$, i.e. at the microchannel exit.

### 5 - Control for caspofungin application

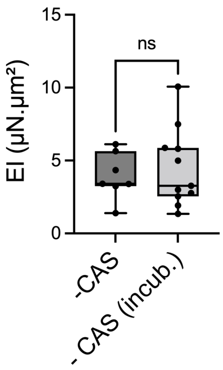


Figure S5 - Influence of the incubation time in the absence of caspofungin on the bending stiffness. The condition -CAS (incub.) correspond to data collected from hyphae submitted to a 30min step in the incubator after an initial measurement (-CAS).

### 6 - Paired data associated to bending stiffness measurements

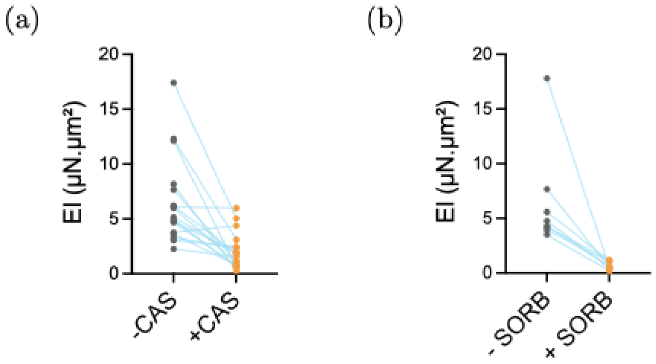


Figure S6 - Distribution of bending stiffness values before and after caspofungine (a) and sorbitol (b) application. Paired data corresponding to figures 4c and 5b right are represented.

### 7 - Influence of hyphae length on EI

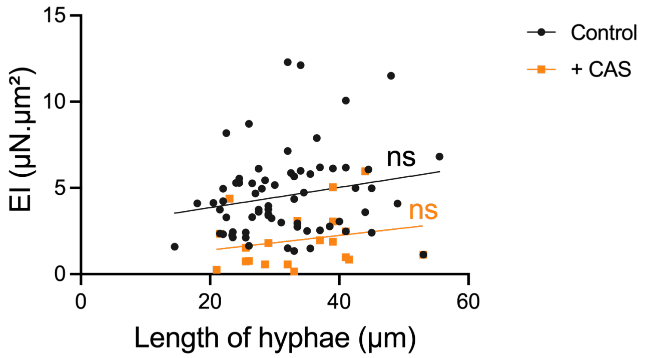


Figure S7 - Bending stiffness as a fonction of hyphal length in SC5314 hyphae without (Control) and after caspofungin application (+CAS). The linear fits show not trend.
